# pSPObooster: a plasmid system to improve sporulation efficiency of *Saccharomyces cerevisiae* lab strains

**DOI:** 10.1101/2024.03.20.586023

**Authors:** Raphael Loll-Krippleber, Yangyang Kate Jiang, Grant W. Brown

## Abstract

Common *S. cerevisiae* lab yeast strains derived from S288C have meiotic defects and therefore are poor sporulators. Here, we developed a plasmid system containing corrected alleles of the *MKT1* and *RME1* genes to rescue the meiotic defects and show that standard BY4741 and BY4742 strains containing the plasmid display faster and more efficient sporulation. The plasmid, pSPObooster, can be maintained as an episome and easily cured or stably integrated into the genome at a single locus. We demonstrate the use of pSPObooster in low- and high-throughput yeast genetic manipulations and show that it can expedite both procedures without impacting strain behavior.

**Take Away:**

- pSPObooster contains corrected alleles or *RME1* and *MKT1*.
- pSPObooster can be maintained as an episome or integrated.
- pSPObooster increases sporulation efficiency by up to 13-fold.
- pSPObooster can be used to speed up high-throughput yeast strain engineering.

## Introduction

The baker’s yeast *Saccharomyces cerevisiae* has been used as a basic and applied research tool for decades in part due to the availability of a large array of genetic and genomic tools. The first *S. cerevisiae* strain with a completely sequenced genome was S288C [Mewes et al. 1997], and as such strains derived from S288C are the workhorses for functional genomics analyses [Mortimer and Johnston 1986; Winston, Dollard, and Ricupero‐Hovasse 1995; Brachmann, Davies, Cost, Caputo, Li, Hieter, and Boeke 1998; Ghaemmaghami, Huh, Bower, Howson, Belle, Dephoure, O’Shea, and Weissman 2003; Huh, Falvo, Gerke, Carroll, Howson, Weissman, and O’Shea 2003; Douglas et al. 2012; Costanzo et al. 2016; Arita et al. 2021]. However, despite their extensive use to study a myriad of biological processes, common yeast lab strains derived from the S288C ancestor including the designer deletion strains BY4741 and BY4742 [Brachmann et al. 1998] are notoriously inefficient at meiosis which can hinder genetic manipulation and analysis as well as limit their use in particular areas of yeast research including population genomics, meiosis, and cell aging, where other yeast backgrounds are preferred.

Genetic defects underlying the poor sporulation efficiency of S288C strains have been identified and mapped to single nucleotide polymorphisms (SNPs) in three genes: *RME1, MKT1*, and *TAO3* [Deutschbauer and Davis 2005]. For *RME1*, which encodes a negative regulator of the meiotic transcriptional program, an adenine deletion at position -308 upstream of the START codon (*rme1-(del308A)*) results in higher gene expression that leads to inhibition of the meiosis process [Bushkin, Pincus, Morgan, Richardson, Lewis, Chan, Bartel, and Fink 2019; Deutschbauer and Davis 2005]. In the case of *TAO3*, which encodes a member of the RAM cell morphogenesis network, a non-synonymous SNP at nucleotide 4477 causes a glutamine to glutamic acid substitution at amino acid 1493 (*tao3-1493QE*), resulting in deregulation of a transcription network important for entry into sporulation [Deutschbauer and Davis 2005; Gupta, Radhakrishnan, Nitin, Raharja-Liu, Lin, Steinmetz, Gagneur, and Sinha 2016]. Finally, a loss-of-function SNP at nucleotide 89 of *MKT1* causes a glycine to aspartate change at amino acid 30 (*mkt1-30D*), impacting a transcription network that regulates the early phases of sporulation [Gupta, Radhakrishnan, Raharja-Liu, Lin, Steinmetz, Gagneur, and Sinha 2015; Deutschbauer and Davis 2005]. Correction of all three alleles to those found in the efficiently-sporulating strain SK1 rescues ∼60% of the S288C sporulation deficiency after 48h of sporulation [Deutschbauer and Davis 2005]. Corrections of *rme1-(del-308A), tao3-1493QE*, and *mkt1-30D* have been engineered together with alleles of four other genes that improve mitochondrial stability and heterologous protein production to yield a new set of lab strains, the DHY strains, which have greatly improved sporulation efficiency [Harvey et al. 2018]. However, because seven allele corrections were introduced, and are not marked, they are not readily crossed into the more standard BY strain derivatives.

In this study we developed a plasmid system to improve the efficiency of sporulation in yeast lab strains derived from S288C. We engineered a plasmid, pSPObooster, carrying the corrected (SK1) alleles of *MKT1* and *RME1* and generated new yeast lab strains containing the plasmid either as an episomal or integrated copy. Our plasmid system significantly accelerates sporulation and increases sporulation efficiency by 13-fold (25% of cells sporulated) after only 3 days in sporulation medium. We also demonstrate the use of pSPObooster to expedite high-throughput strain engineering and genetic screens that use Synthetic Genetic Array technology.

## Material and methods

### Yeast maintenance and growth conditions

Strains used in this study are listed in Table 1. Yeast strains were grown at 30°C in liquid or solid standard rich (YPD; 20 g/L peptone, 20g/L dextrose, 10 g/L yeast extract and 20 g/L agar for solid medium) or synthetic (SD; 6.7 g/L yeast nitrogen base, 20 g/L glucose and 20 g/L agar for solid medium) medium containing all amino acids minus the relevant amino acids when auxotrophic marker selection was needed [Dunham, Gartenberg, and Brown 2015]. To induce sporulation, yeasts were cultured in liquid sporulation medium (SPO: 1% potassium acetate and all amino acids at 0.25 times the concentration in synthetic medium) at 25°C.

**Table 1.**
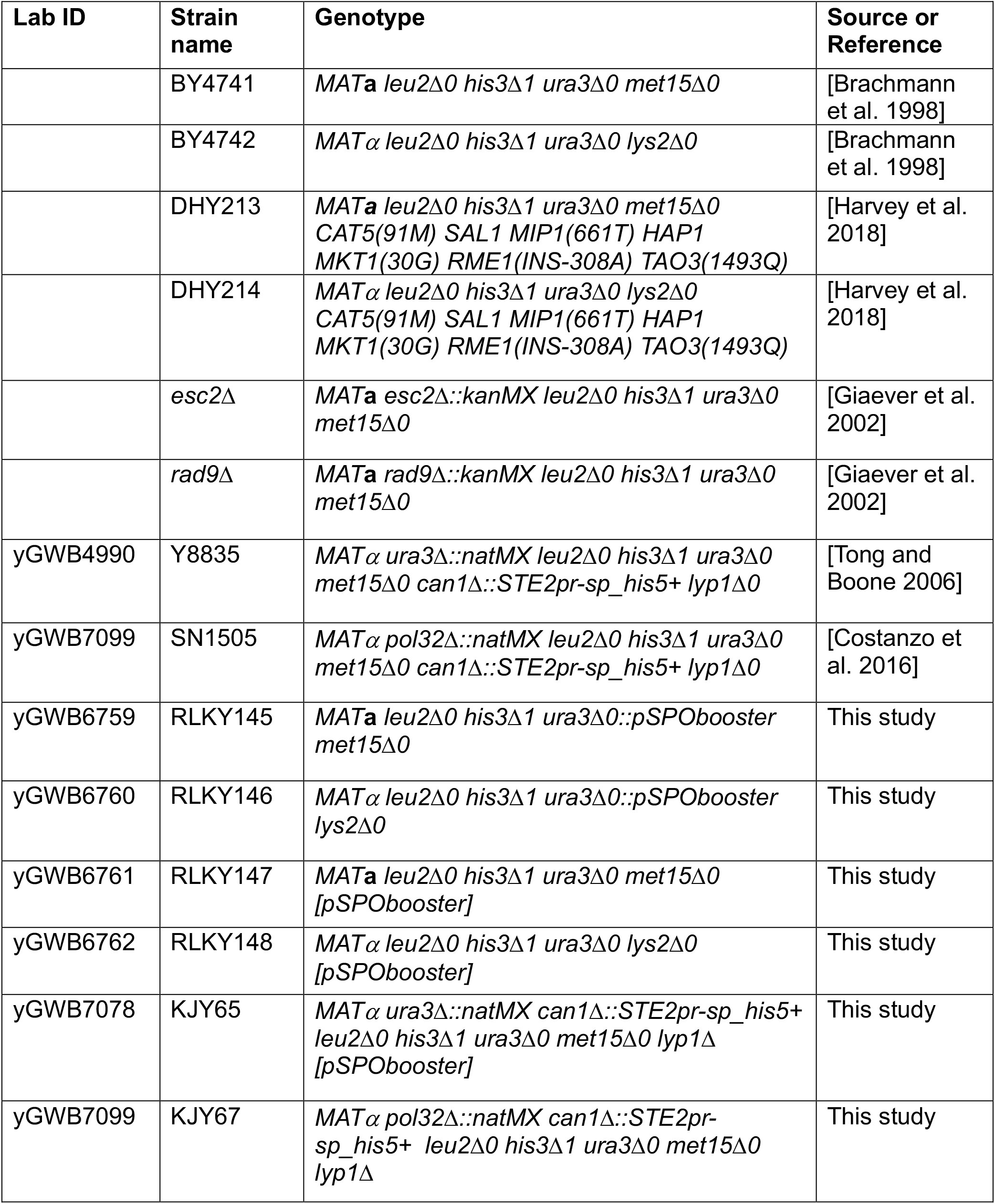
Strains used in this study.

| Lab ID | Strain name | Genotype | Source or Reference |
| --- | --- | --- | --- |
|  | BY4741 | *MATa leu2Δ0 his3Δ1 ura3Δ0 met15Δ0* | [Brachmann et al. 1998] |
|  | BY4742 | *MATα leu2Δ0 his3Δ1 ura3Δ0 lys2Δ0* | [Brachmann et al. 1998] |
|  | DHY213 | *MATa leu2Δ0 his3Δ1 ura3Δ0 met15Δ0 CAT5(91M) SAL1 MIP1(661T) HAP1 MKT1(30G) RME1(INS-308A) TAO3(1493Q)* | [Harvey et al. 2018] |
|  | DHY214 | *MATα leu2Δ0 his3Δ1 ura3Δ0 lys2Δ0 CAT5(91M) SAL1 MIP1(661T) HAP1 MKT1(30G) RME1(INS-308A) TAO3(1493Q)* | [Harvey et al. 2018] |
|  | *esc2Δ* | *MATa esc2Δ::kanMX leu2Δ0 his3Δ1 ura3Δ0 met15Δ0* | [Giaever et al. 2002] |
|  | *rad9Δ* | *MATa rad9Δ::kanMX leu2Δ0 his3Δ1 ura3Δ0 met15Δ0* | [Giaever et al. 2002] |
| yGWB4990 | Y8835 | *MATα ura3Δ::natMX leu2Δ0 his3Δ1 ura3Δ0 met15Δ0 can1Δ::STE2pr-sp_his5+ lyp1Δ0* | [Tong and Boone 2006] |
| yGWB7099 | SN1505 | *MATα pol32Δ::natMX leu2Δ0 his3Δ1 ura3Δ0 met15Δ0 can1Δ::STE2pr-sp_his5+ lyp1Δ0* | [Costanzo et al. 2016] |
| yGWB6759 | RLKY145 | *MATa leu2Δ0 his3Δ1 ura3Δ0::pSPObooster met15Δ0* | This study |
| yGWB6760 | RLKY146 | *MATα leu2Δ0 his3Δ1 ura3Δ0::pSPObooster lys2Δ0* | This study |
| yGWB6761 | RLKY147 | *MATa leu2Δ0 his3Δ1 ura3Δ0 met15Δ0 [pSPObooster]* | This study |
| yGWB6762 | RLKY148 | *MATα leu2Δ0 his3Δ1 ura3Δ0 lys2Δ0 [pSPObooster]* | This study |
| yGWB7078 | KJY65 | *MATα ura3Δ::natMX can1Δ::STE2pr-sp_his5+ leu2Δ0 his3Δ1 ura3Δ0 met15Δ0 lyp1Δ [pSPObooster]* | This study |
| yGWB7099 | KJY67 | *MATα pol32Δ::natMX can1Δ::STE2pr-sp_his5+ leu2Δ0 his3Δ1 ura3Δ0 met15Δ0 lyp1Δ* | This study |

For the synthetic genetic array diploid selection, the following media were used: YPD+G418+clonNAT (YPD containing 200 µg/mL G418 and 100 µg/mL clonNAT respectively) or SD/MSG-ura+G418+clonNAT (1.7 g/L YNB without nitrogen source, 1 g/L monosodium glutamate, 20 g/L glucose, 20 g/L agar, 200 µg/mL G418, 100 µg/mL cloNAT) for diploid selection. For the SGA sporulation, haploid and double mutant selection steps, enriched sporulation medium, SD/MSG-his-lys-arg+CAN+SAEC and SD/MSG-his-lys-arg+CAN+SAEC+G418+clonNAT were used, respectively, and prepared as described in [Tong and Boone 2006].

### Plasmid construction

The pSPObooster plasmid was constructed using yeast plasmid assembly. First, the *mkt1-30G* and *rme1-ins308A* alleles (including ∼1000 bp upstream and ∼ 200bp downstream regulatory sequences) were amplified from DHY214 [Harvey et al. 2018] genomic DNA using RLO336/RLO338 and RLO337/RLO339 primers (Table 2) which introduce sequences homologous to the regions flanking the *Sac*I and *Hind*III sites of pRG216 (Addgene #64528) [Gnügge and Rudolf 2017]. Plasmid assembly was performed by transforming yeast strain BY4741 with 100 ng of *Sac*I/*Hind*III linearized pRG216 along with the *mkt1-30G* and *rme1-ins308A* PCR products, and selecting for Ura+ transformants. Correct plasmid assembly was verified by whole plasmid sequencing following plasmid recovery into *Escherichia coli*. The plasmid sequence is provided in the Supplemental Material and pSPObooster is available from Addgene (#216160).

**Table 2.**
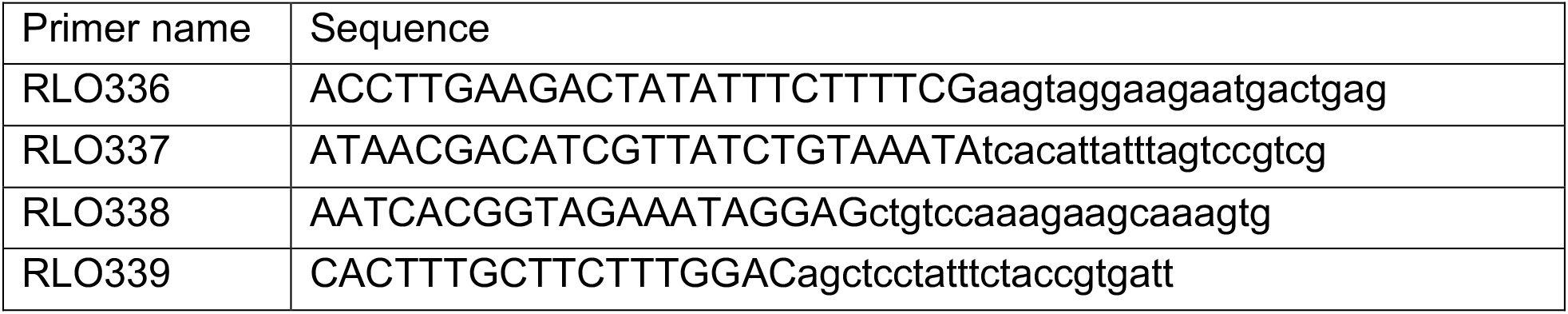
Primers used in this study. Uppercase letters represent homology introduced for yeast plasmid assembly into pRG216.

### Yeast transformation and strain construction

Yeasts were transformed using the standard lithium acetate transformation procedure [Dunham, Gartenberg, and Brown 2015]. BY4741 or BY4742 were transformed with circular pSPObooster to yield RLKY147 and RLKY148, or with *Asc*I-linearized pSPObooster to integrate the *mkt1-30G* and *rme1-ins308A* genes at the *ura3Δ0* locus, yielding RLKY145 and RLKY146 (Table 1). Correct integrations were confirmed by PCR. Synthetic Genetic Array (SGA) strains were constructed by transforming pSPObooster into *ura3Δ* (Y8835) or *pol32Δ* (SN1505) BY4742 SGA query strains to yield KJY65 and KJY67 (Table 1).

### Sporulation efficiency

Haploid cells of opposite mating type were mated on solid YPD for 24 hours before streaking cells on SD-met-lys or SD-met-lys-ura (if plasmid selection was required) to obtain single diploid colonies. A single diploid colony was used to inoculate 5 mL of liquid SD with all amino acids or SD-ura (if plasmid selection was required) and grown at 30°C overnight. One mL of overnight culture was harvested by centrifugation, 900 µl of supernatant was removed, and the remaining 100 µl was resuspended in 5 mL of sporulation medium and incubated at 23°C for up to 4 days. Cells from the sporulation culture were sampled at regular times, fixed with 70% ethanol, and imaged on a Zeiss AxioImager Z1 microscope equipped with a 63X brightfield objective. Sporulation efficiency was assessed by counting the numbers of cells and asci in the cultures at each timepoint, dividing the number of asci by the number of cells plus asci, and multiplying by 100. At least 100 cells/asci were counted for each sample.

### Synthetic Genetic Array procedure

SGA analysis was performed as previously described [Tong and Boone 2006], with the following modifications: Using a Biomatrix colony pinning robot (S&P Robotics), strains KJY65 and KJY67 were mated with one 1536 colony array plate carrying 384 gene deletion mutant strains each in quadruplicate (plate 3 of the SGAv2 array) on YPD and SD-ura for two days at room temperature. Diploids were selected by pinning onto SD/MSG-ura+G418+clonNAT and growing for two days at 30°C. Diploid cells were then pinned onto three SPO plates and were incubated for 24h, 48h, and 72h at 23°C. Spores were then pinned on haploid selection media and imaged after two days to assess sporulation efficiency. In parallel, the untransformed *ura3Δ* (Y8835) and *pol32Δ* (SN1505) query strains were subjected to the same SGA selection scheme, except that mating was performed on YPD and diploid selection on YPD+G418+clonNAT. Diploid cells were then pinned onto four sporulation plates and incubated for 24h, 48h, 72h, or 7 days at 23°C. Spores were then pinned on haploid selection media and imaged after two days to assess sporulation efficiency.

After imaging the haploid selection plates, the haploid strains obtained after 72h of sporulation for KJY65 and KJY67, and after 7 days of sporulation for Y8835 and SN1505, were taken through the rest of the SGA procedure, ultimately pinning to media that selects for haploid double mutants to allow scoring for synthetic genetic interactions. Colony sizes were measured and genetic interactions scored using SGAtools [Wagih et al. 2013] after two days of growth at 30°C on double mutant selection medium (Tables S2 and S3). Negative genetic interactions with an SGA score < -0.12 and p-value < 0.05 were compared to *pol32*Δ negative genetic interactions meeting the same criteria, obtained from the CellMap [Costanzo et al. 2016] (Table S1)

### High-throughput sporulation efficiency

Colony sizes measured on SGA haploid selection medium after sporulation were used as an indirect proxy for sporulation efficiency. Colony size was determined after 2 days of growth at 30°C on haploid selection medium using the gitter package [Wagih and Parts 2014]. Border colonies, which are large due to a spatial artifact associated with nutrient supply, were removed from the analysis and raw colony size from the remaining colonies was plotted using a custom pipeline in R (https://www.r-project.org/).

### Plasmid Loss Assessment

A control colony (*his3*Δ *ura3*Δ or *his3*Δ *pol32*Δ from the KJY65 or KJY67 crosses) from an SGA double mutant selection plate was resuspended in sterile water and cell concentration was determined with a Z1 Beckman-Coulter particle counter equipped with a 100 µm aperture tube. Cells were diluted to obtain single colonies after plating 200 µL of cell suspension on either YPD or SD-ura. Plates were incubated for two days at 30°C before assessing the number of Ura+ colonies relative to the total colony number obtained on YPD.

## Results and discussion

### Design of a plasmid system to boost sporulation efficiency

We set out to engineer a system allowing easy correction of the sporulation defect observed in S288C derived lab strains. To this end, we designed a low copy CEN-ARS plasmid based on the pRG216 backbone [Gnügge and Rudolf 2017] and containing the *mkt1-30G* and *rme1-ins308A* alleles [Deutschbauer and Davis 2005] (Figure 1A). Due to plasmid size considerations, the *tao3*-*1493Q* allele was omitted from the design as the *TAO3* gene with its regulatory regions is approximately 8500bp. Although *tao3*-*1493Q, mkt1-30G* and *rme1-ins308A* together gave the most effective increase in sporulation, the combination of *mkt1-30G* and *rme1-ins308A* alone was sufficient to increase sporulation by ∼5-fold, to ∼40% of the efficiently sporulating SK1 strain [Deutschbauer and Davis 2005]. Therefore, we reasoned that the improvement offered by combining *mkt1-30G* and *rme1-ins308A* would be sufficient for routine genetic studies and manipulations. The resulting plasmid, pSPObooster, can be selected using the *URA3* marker and counter-selected on 5-FOA, and maintained as an episome or integrated at the *ura3*Δ*0* locus after digestion with the *Asc*I restriction enzyme (Figure 1A and B).

**Figure 1.**
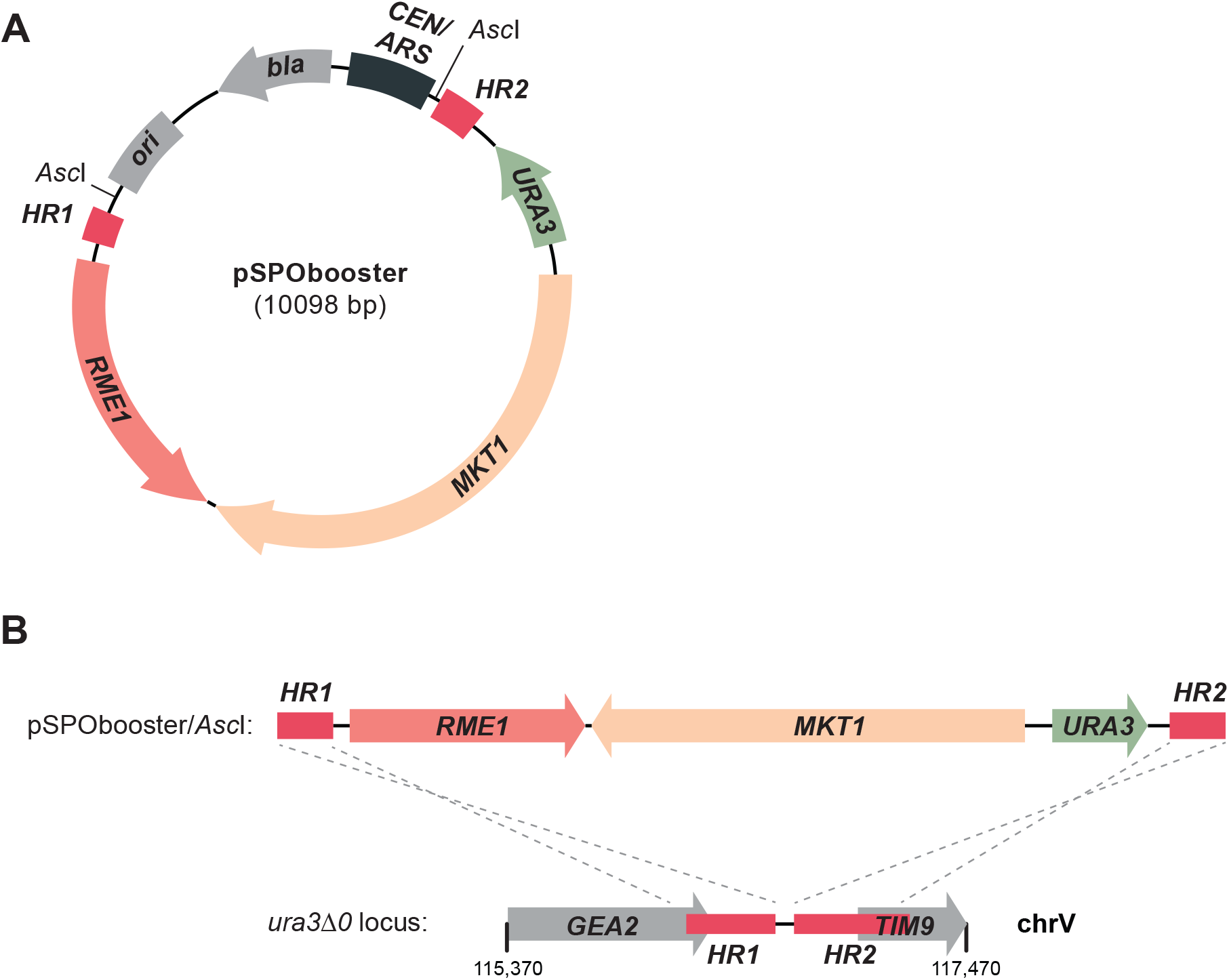
Schematic map of the sporulation booster plasmid. **A**. Map of pSPObooster. Genetic elements and their orientations are depicted as colored arrows. *MKT1* and *RME1* were amplified from DHY214 and cloned into pRG216. **B**. Map of pSPObooster integrated at the *ura3Δ0* locus after *Asc*I restriction digestion. Genetic elements and their orientation are depicted as colored arrows. The *ura3*Δ*0* locus lacks the *URA3* open reading frame and the 5’ and 3’ untranslated regions. pSPObooster integration occurs via homologous recombination between the HR1 and HR2 regions in pSPObooster and their corresponding sequences upstream and downstream of the *URA3* locus.

### Efficient and rapid sporulation is achieved using the sporulation booster plasmid

In order to test whether pSPObooster can restore high sporulation efficiency, BY4741 and BY4742 (strains derived from S288C) were transformed to introduce pSPObooster as an episome or single-copy integrant, and mated with a non-transformed BY strain of the opposite mating type to generate diploids. A highly efficient sporulating control strain was constructed using *MAT***a** and *MAT*α cells of the DHY strain background carrying corrected (SK1) alleles of *MKT1, RME1* and *TAO3* at their cognate loci [Harvey et al. 2018]. After 48h of sporulation in liquid medium cells carrying pSPObooster either as an episome or as an integrated copy were 2.1% and 1.4% sporulated, respectively (Figure 2). No asci were detected with BY diploids, and 50% of cells had sporulated in the DHY diploid culture. After 72h of sporulation, the strains containing pSPObooster plasmid were 24% sporulated while BY diploids displayed only 2% sporulation and DHY diploids reached 68% sporulation. After 4 days, the strains with pSPObooster plasmid reached 50% sporulation, DHY diploids were 81% sporulated, and the BY diploids were only 14% sporulated.

**Figure 2.**
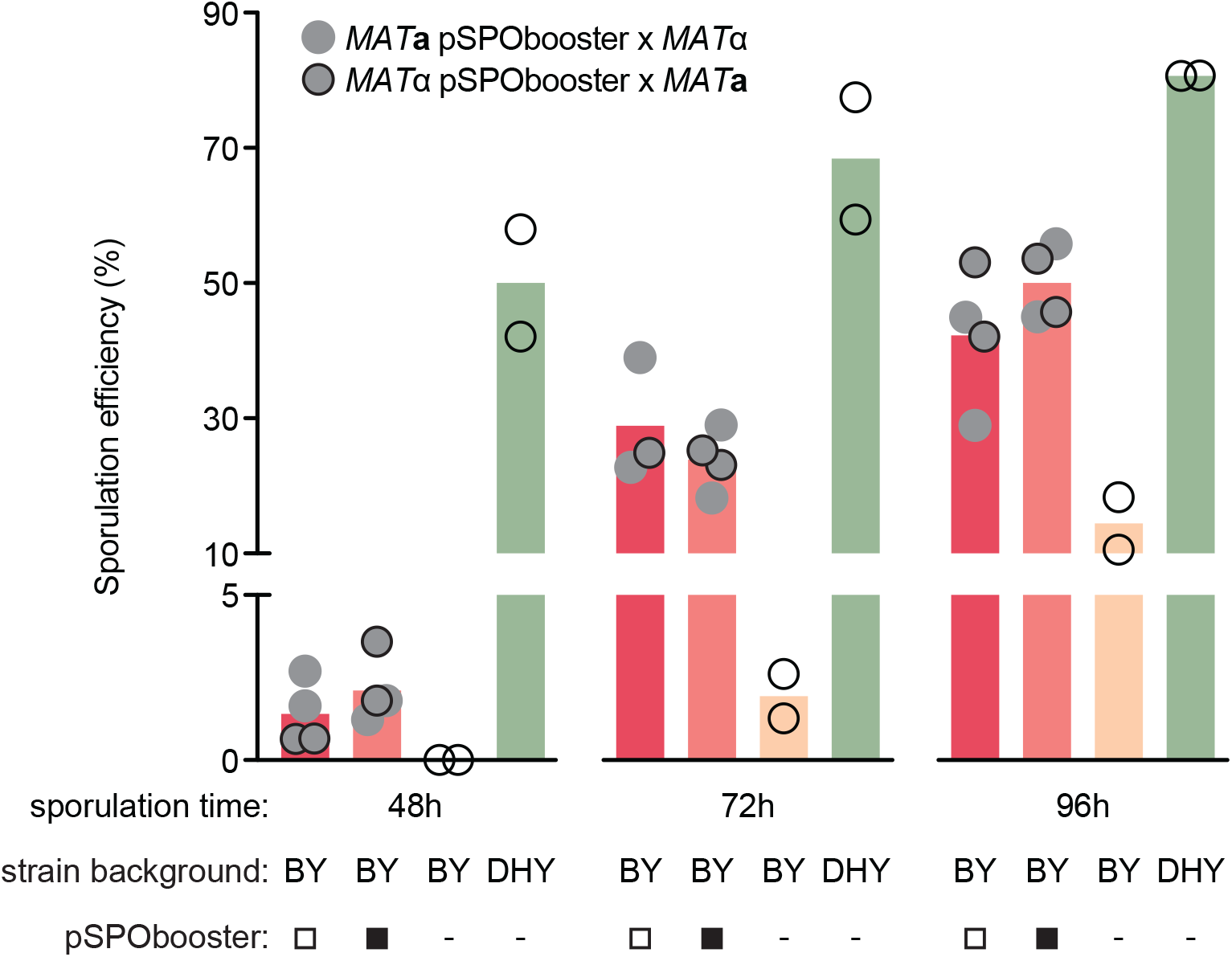
The sporulation booster plasmid is functional and promotes efficient sporulation in 3 days. The sporulation efficiencies of the indicated diploid strains are plotted. *MAT***a** or *MATα* haploid strains carrying pSPObooster as an integrated copy (RLKY145 and RLKY146, open squares) or as an episome (RLKY147 and RLKY148, closed squares) were mated with BY strains of the opposite mating type to generate diploids. Diploids were sporulated and SPO cultures were sampled at the indicated timepoints. Untransformed BY (BY4741 x BY4742) and DHY (DHY213 x DHY214) diploids were included as inefficient and efficient sporulation controls, respectively. At least 100 cells were counted for each sample. n = 2.

We conclude that pSPObooster is functional and promotes faster and more efficient sporulation of the S288C-derived BY strains. As expected, diploids with pSPObooster sporulated less efficiently than DHY diploids, likely due to the additional *TAO3* correction in DHY strains.

### Fast and efficient high-throughput genetic manipulation using pSPObooster

Next tested if pSPObooster can accelerate high-throughput genetic engineering using Synthetic Genetic Array (SGA) technology [Tong and Boone 2006]. The *POL32* gene, which encodes a subunit of DNA polymerase δ, has been widely characterized and has a rich genetic interaction network [Costanzo et al. 2016], making it a good test case for pSPObooster. We transformed *pol32*Δ and *ura3*Δ BY4742 SGA query strains with pSPObooster and performed SGA with one 1536 colony plate of the non-essential gene deletion collection colony array. We included the untransformed *pol32*Δ and *ura3*Δ BY4742 query strains as controls. We selected the colony array plate with the highest number of genetic interactions with *POL32* according to the CellMap [Costanzo et al. 2016]. The SGA query strains were mated with the array on rich (YPD) or minimal (SD-ura) media, replica pinned to sporulation medium after diploid selection, and allowed to sporulate for 24h to 7 days.

After sporulation, colonies were pinned to media that selects for haploids. We reasoned that colony size after haploid selection should be a good proxy for sporulation efficiency, as more spores will yield more haploid cells and therefore larger colonies when haploids are selected. We measured colony sizes after 2 days growth on haploid selection medium for the 24h, 48h, 72h and 7-day sporulations (Figure 3A). Both the *ura3*Δ and *pol32*Δ strains carrying pSPObooster gave rise to bigger haploid colonies at all sporulation times (Figure 3A, compare green to orange boxes at each time). The average colony size for BY strains after 7 days of sporulation (the standard sporulation time for SGA) was smaller than that obtained with pSPObooster after 72h of sporulation, indicating that use of pSPObooster can reduce the sporulation time in SGA protocols by more than half.

**Figure 3.**
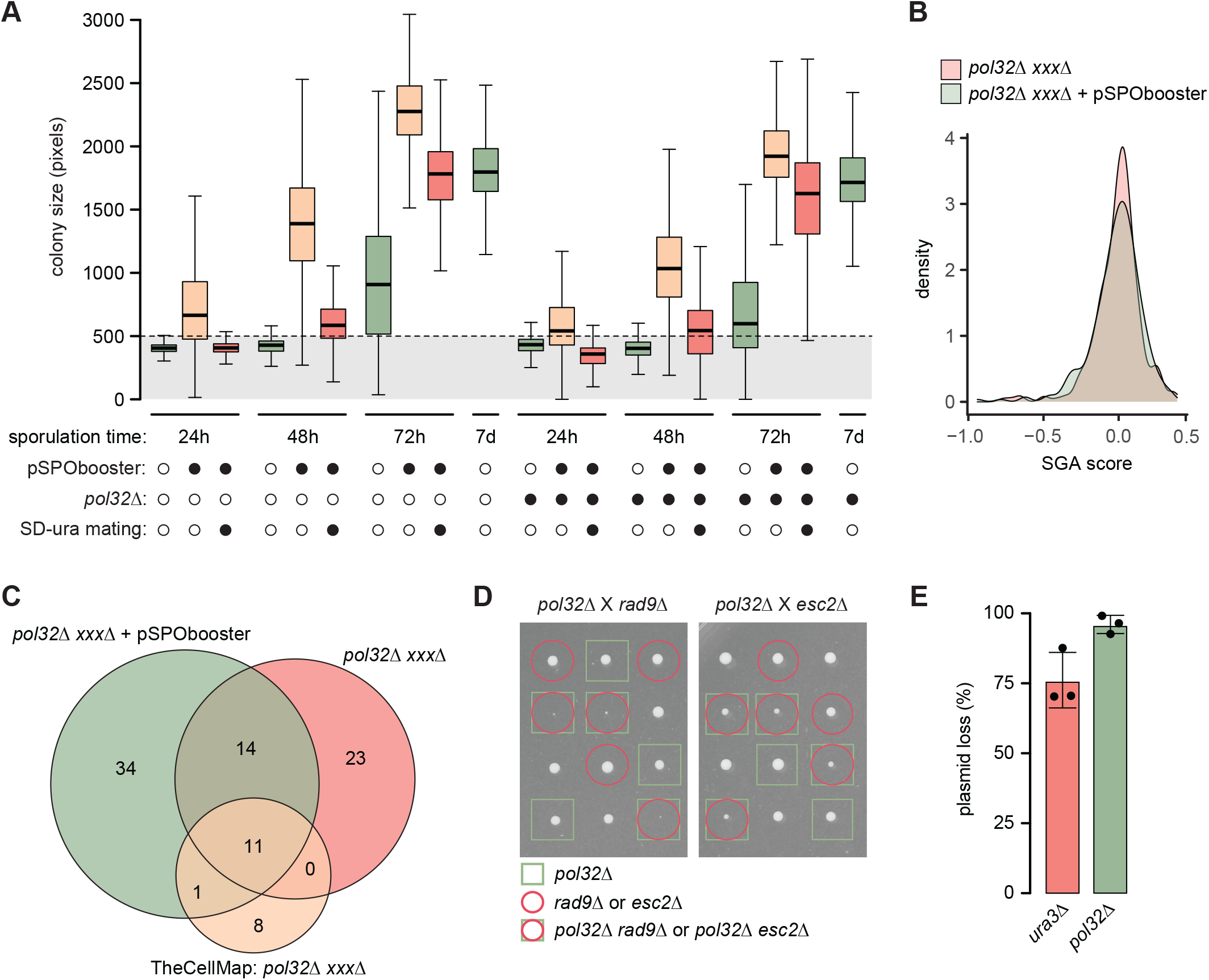
The pSPObooster plasmid allows rapid and efficient high-throughput genetic crosses. **A**. Colony sizes on haploid selection medium after sporulation for 24h, 48h, 72h, or 7 days are shown as boxplots for the indicated strains. A *ura3Δ;* (open circles) or *pol32Δ* (closed circles) strain was mated to a 1536 colony array of yeast gene deletion mutants. The *ura3*Δ and *pol32*Δ strains were either untransformed (open circles) or carried pSPObooster as an episome (closed circles) and were mated on YPD (open circles) or SD-ura (closed circles). Boxes span the first to third quartile values, the medians are indicated by the horizontal black bars, and the whiskers span the minimum to maximum values. The horizontal dotted line represents the lowest acceptable haploid colony size (500 pixels) in the analysis. 1232 colonies were measured. **B**. SGA score distributions for *pol32Δ xxxΔ* double mutants with or without pSPObooster. **C**. Venn diagram comparing the genetic interactions identified in the *pol32*Δ SGA screens with or without pSPObooster and *POL32* genetic interactions annotated in the TheCellMap. **D**. Tetrad analysis of crosses between a *pol32*Δ strain carrying pSPObooster and either *rad9*Δ or *esc2*Δ strains. Each column is the four haploid segregants from a single tetrad. The genotype of each segregant is indicated. **E**. The fraction of cells that had lost pSPObooster at the end of the SGA procedure is plotted for haploid double mutants from the control SGA (*ura3*Δ *his3*Δ) and the *POL32* SGA (*pol32*Δ *his3*Δ). n=3.

ARS/CEN plasmids like pSPObooster are readily lost in the absence of selection [Zhang, Moo-Young, and Chisti 1996], and so we tested whether plasmid selection during the mating step is important for optimal sporulation in a high-density colony array. We measured colony sizes on haploid selection medium after colony array matings on SD-ura (Figure 3A, red boxes). We found that cells mated on YPD gave bigger haploid colony sizes compared to cells mated on SD-ura for all strains for all sporulation times. We conclude that plasmid selection during mating is not important for the efficient sporulation conferred by pSPObooster.

To test whether pSPObooster is compatible with high-throughput genetic interaction screening, we proceeded with the double-mutant selection steps of the SGA using the pSPObooster haploid strains after 72h of sporulation and the untransformed haploid strains after 7 days of sporulation. The distributions of genetic interaction scores were very similar with and without pSPObooster (Figure 3B). Considering high-confidence genetic interactions (SGA scores < -0.12 with p-value < 0.05, [Costanzo et al. 2016]), the SGA screen with pSPObooster yielded 60 interactions, of which 25 (42%) overlapped with the SGA screen without the plasmid. The pSPObooster screen recovered 12 of 20 the known *POL32* interactions, while the control screen recovered 11 of 20 (Figure 3C) [Costanzo et al. 2016]. Interestingly, the screen with pSPObooster gave the highest number of genetic interactions. Considering that the plasmid gives very robust growth at the haploid selection step after 72 hours of sporulation, even as compared to 7 days of sporulation for the untransformed strains, we hypothesize that the shorter sporulation time could result in increased dynamic range to allow detection of more subtle fitness defects after double mutant selection.

To further demonstrate that pSPObooster did not impact the detection of genetic interactions, we performed tetrad analysis on *pol32Δ* pSPObooster x *rad9Δ and polΔ32* pSPObooster x *esc2Δ* crosses. Genetic interactions between *POL32* and *RAD9*, and *POL32* and *ESC2* were readily observable (Figure 3D), confirming that pSPObooster does not impair detection of known *pol32*Δ genetic interactions. Finally, we assessed the proportion of cells retaining pSPObooster at the end of the SGA procedure. Between 75% (*ura3*Δ) and 90% (*pol32*Δ) of cells had lost the plasmid (Figure 3E), suggesting that the plasmid is unlikely to have major impact on genetic interactions scored at the end of the SGA procedure. Nevertheless, because pSPObooster contains a counter-selectable marker, plasmid curing before double mutant selection and scoring is readily achievable using 5-FOA.

## Conclusion

We developed a plasmid system that allows robust improvement of the sporulation defect observed in one of the most widely used *S. cerevisiae* lab strain backgrounds, S288C. Because our system is plasmid-based and counter-selectable, low sporulation efficiency can be easily corrected in any S288C-derived strain using a simple plasmid transformation. Alternatively, the sporulation booster plasmid can be readily integrated at the *URA3* locus, in *ura3* and *ura3Δ0* strains, allowing stable retention of the efficient sporulation phenotype while being easily trackable. To illustrate the benefits of our system, we used pSPObooster in a high-throughput yeast genetics experiment (SGA) and showed that not only can it accelerate the entire procedure but also recapitulates established genetic interactions. We anticipate that the use of pSPObooster will facilitate routine genetic manipulations and high-throughput genetics in yeast strains derived from S288C.

## Acknowledgements.

This work was supported by Natural Sciences and Engineering Research Council of Canada grant RGPIN-2017-06855 to GWB. GWB holds a Tier 1 Canada Research Chair. We are grateful to work on the lands of the Mississaugas of the Credit, the Anishnaabeg, the Haudenosaunee and the Wendat peoples, land that is now home to many diverse First Nations, Inuit, and Métis peoples.

## Conflict of Interest Statement

The authors declare no conflict of interest.

## Notes

### Competing Interest Statement

The authors have declared no competing interest.

